# Differential effects of emerging broad-spectrum streetlight technologies on landscape-scale bat activity

**DOI:** 10.1101/525568

**Authors:** Rowland Williams, Charlotte Walters, Rory Gibb, Ella Browning, David Tipping, Kate E. Jones

**Affiliations:** Centre for Biodiversity and Environment Research, Department of Genetics, Evolution and Environment, University College London, Gower Street, London, WC1E 6BT, United Kingdom; Institute of Zoology, Zoological Society of London, Regent’s Park, London, NW1 4RY, United Kingdom; Department of the Environment, States of Jersey, Howard Davis Farm, La Route de la Trinité, Trinity, JE3 5JP, Jersey, Channel Islands

**Author notes:** Corresponding authors: Rowland Williams; Kate E. Jones.

**Keywords:** bats, acoustic monitoring, artificial lighting, citizen science, island, roads, States of Jersey, urbanisation

## Abstract

Urbanization has greatly reduced the extent of high quality habitat available to wildlife with detrimental consequences documented across a range of taxa. Roads and artificial lighting regimes are dominant features of the modern environment, and there is currently a rapid worldwide transition towards energy-efficient, broad-spectrum white-light streetlight technologies such as metal halide (MH) and more recently, light-emitting diode (LED), despite little being known about their broad ecological impacts. Here, in a five-year citizen science study across the island of Jersey, we combine detailed lighting and habitat data with ultrasonic bat survey data collected from 2011 to 2015 (before and after a LED lighting technology transition), to analyse the landscape-scale effects of different broad-spectrum streetlight technologies on activity of a widespread, generalist bat species. In contrast to many experimental studies, we show that the local density of both traditional yellow high-pressure sodium (HPS) and more modern LED streetlights have significant negative effects on activity of the common pipistrelle *(Pipistrellus pipistrellus)* compared to unlit areas, while accounting for spatial bias, bat population trends over time, surrounding habitat type and road-type. In contrast, we find no discernable impact of the density of ultra-violet emitting MH lighting on bat activity. This is the first large-scale evidence that emerging artificial lighting technologies have differential impacts on activity, even for a bat species generally characterised as light-tolerant and commonly found in urban areas. Importantly, our landscape-level approach also demonstrates that the degree of urbanization and road type have even larger negative impacts on bat activity, independent of artificial lighting regime. Our findings emphasise the need for improving landscape-scale understanding of the ecological impacts of new lighting technologies prior to widespread uptake, and have important implications for future streetlight installation programmes and urban planning more generally.

## Introduction

Global biodiversity is declining at unprecedented rates in response to anthropogenic pressures (1). Urbanization has greatly reduced the extent of high quality habitat available to biodiversity (2), and roads in particular have caused widespread fragmentation of landscapes (3), creating barriers to animal movement, severing commuting routes and restricting access to foraging sites (4). A key feature of modern road environments is artificial light at night (ALAN). ALAN levels are increasing rapidly; from 2012 to 2016, the global artificially lit outdoor area increased by 2.2% per year (5). ALAN is considered an emerging threat to biodiversity, as detrimental effects have been documented in a range of taxa. For example, ALAN may disrupt networks of nocturnal pollinators which could have consequences for the provision of important ecosystem services (6). Even low levels of artificial light may be capable of causing significant phenological shifts, such as early onset of reproduction in songbirds (7). Artificial lighting regimes may therefore exacerbate the barrier effects of roads on wildlife by increasing energetic costs and reducing long-term fitness (8).

Bats (order Chiroptera) are of conservation importance as almost one quarter of species are threatened globally (9). Bats provide key ecosystem services and are considered an effective bio-indicator across a range of spatial scales (10). Bats are vulnerable to the fragmenting”effects of roads as their home range sizes tend to be larger than would be predicted from body size (11,12). The nocturnal activity patterns of bats make them sensitive to ALAN, and clutter-adapted genera such as *Myotis* and *Rhinolophus* are known to be particularly light-averse. Lighting may force bats to use inferior commuting routes, which may increase energetic costs and predation risk (13). However, fast-flying bat species adapted to foraging in open areas may be relatively light-tolerant and could benefit from aggregations of insect prey around streetlights. Indeed, *Pipistrellus, Nyctalus* and *Eptesicus* bat species have all been observed foraging on insects at streetlights (14–16). However, by creating a ‘vacuum effect’, whereby insect biomass is attracted away from unlit areas, artificial lighting may reduce food availability for light-averse species (17). The presence of streetlights in urban or semi-urban landscapes therefore has the potential to alter the composition of bat assemblages and exacerbate the detrimental impacts of roads on bat ecology (18).

Traditional methods of road illumination involve orange low pressure sodium (LPS) and yellow high pressure sodium (HPS) lamps (19). Recently, there have been widespread transitions toward white-light technologies such as metal halide (MH) and energy-efficient light emitting diode (LED) to reduce operating costs and carbon emissions (20). The ecological consequences of such changes remain unclear. The four technologies can be divided into three categories reflecting spectral composition: non-ultraviolet (UV) emitting narrow spectrum (LPS), non-UV emitting broad spectrum (HPS and LED) and UV emitting broad spectrum (MH) (Table 1) (19). Experimental evidence suggests that the effects of different streetlight technologies on bat behaviour may be dependent on ultraviolet (UV) content (18). In general, bat activity for more light-tolerant species is positively associated with UV content, probably due to increased attraction of insect prey (17,21).

**Table 1:** Spectral bandwidth and UV-content of the four main streetlight types. In our study area, only HPS, LED and MH lights were present.

| Light-type | Spectral bandwidth | UV-emission |
| --- | --- | --- |
| Low pressure sodium (LPS) | Narrow | No |
| High pressure sodium (HPS) | Broad | No |
| Light-emitting diode (LED) | Broad | No |
| Metal halide (MH) | Broad | Yes |

Given the interspecific variation in light-tolerance among bats, the effects of different lighting technologies are also likely to be species-specific. For example, experimental illumination of hedgerows with HPS and LED lights led to a significant reduction in *Rhinolophus hipposideros* activity but had no effect on *Pipistrellus* sp. activity (8,22). However, other work at larger spatial scales has shown that *Pipistrellus* activity is reduced in road sections lit with HPS lights (23), suggesting that the observed effects of different light types may be dependent on the spatial scale at which analyses are conducted. Although at the landscape scale there appears to be a broadly negative relationship between ALAN and bat activity (24), the relative effects of different forms of artificial lighting on bat ecology are unknown. Given the current global transition towards white light technologies (21), improved understanding of their impacts is vital to inform effective road mitigation and bat conservation.

Here, we collect acoustic data from roadside transects across the island of Jersey, as part of the Indicator Bats Programme (iBats), a citizen science bat population monitoring scheme (25,26). Using activity data for the common pipistrelle bat, *Pipistrellus pipistrellus* and streetlight data from 2011 to 2015, a period spanning the recent transition from older HPS lights to higher-efficiency LED lights across parts of the island, we analyse the landscape effects of three broad-spectrum streetlight technologies HPS, MH and LED on bat activity, accounting for spatial bias, time (multi-year bat population trends), habitat heterogeneity, climate and road type. We focus on *P.pipistrellus* because roadside acoustic monitoring is biased towards this fast-flying species and, although generally considered to be tolerant of lighting and often found in urban areas, it remains unclear whether ALAN regimes have large-scale effects on this species’ behaviour and ecology (26). Considering this, and the link between UV content and bat activity around streetlights, we expected *Pipistrellus pipistrellus* activity to be lower along roads lit with non-UV emitting HPS and LED lights than those lit with UV emitting MH lights. We also investigate the relative impact of habitat and environmental drivers on bat activity and predict landscape suitability at an island-wide scale.

## Results

During roadside surveys across Jersey from 2011 to 2015, we recorded 2974 bat passes (discrete sequences of calls emitted by an individual) from ten different bat species or species groups, with the most abundant species being the common pipistrelle (*P. pipistrellus*) and least abundant, *Myotis* species (Figures 1b & 2, Supplementary Information Table S2). Records of juvenile *P. kuhlii* have been recently identified on Jersey, making this likely to be a resident species (pers comm. David Tipping 2015), rather than a misclassification of other resident pipistrelle species. We modelled the relationship between *P. pipistrellus* activity and density of street-lighting across sampled 50m^2^ grid-cells using Poisson generalised linear mixed-effects regression, controlling for space, time (multi-year bat population trends), habitat heterogeneity, climate and road type, with model averaging carried out in an information-theoretic framework (see Materials & Methods). Both the local density of HPS and LED lights had a significant and similar negative effect on *P. pipistrellus* activity, whilst the effect of MH density was neutral and insignificant (confidence interval includes zero and cumulative AICc weight (W) is low (<0.4)) (Table 2, Figure 3). The percentage cover of woodland and water had a significant positive effect on *P. pipistrellus* activity whilst urban percentage cover had a significant negative effect (Figure 3). Since all habitat and lighting variables were centred and scaled prior to modelling, our results show that the relative effect of urban cover on bat activity was stronger than all lighting technologies tested (Figure 3). The percentage cover of arable land and grassland around roads were not informative predictors of *P.pipistrellus* activity, as the confidence intervals of these estimates included zero and their cumulative AICc weights were low (<0.6 and <0.4, respectively) (27). Bat activity was also related to the type of road from which bat passes were detected, with the one major road in Jersey having the strongest significant negative impact on *P. pipistrellus* activity followed by main roads (Table 2). Temperature had no significant effect on *P.pipistrellus* activity whereas wind speed had a significant positive effect (Table 2). The spatial autocovariate (see Materials & Methods) was an important predictor variable (cumulative AICc weight of 1) in the average model, and inclusion of the autocovariate in the global model also noticeably improved model fit in terms of AIC, indicating that there was significant autocorrelation in the spatial distribution of sampled bat activity across Jersey.

**Figure 1.**
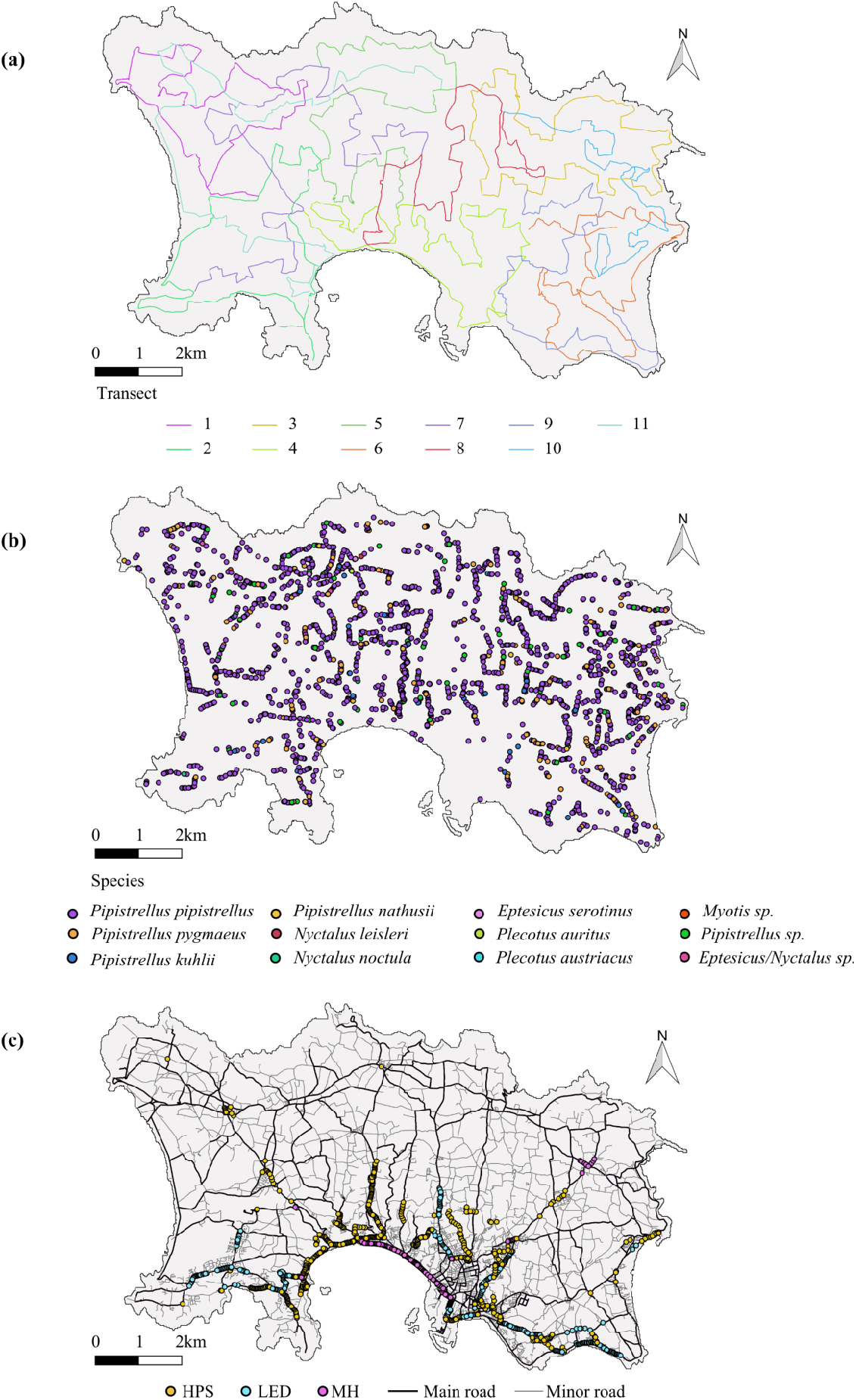
Island of Jersey showing the distribution of (A) transects, (B) roads, road type, and street lighting along road transects, and (C) bat species/species groups recorded. Coloured lines in (A) represent acoustic iBats road transects (n=11), each driven twice yearly from 2011-2015. Different coloured dots in (B) represent different lighting technologies, where yellow is high pressure sodium (HPS), blue is light emitting diode (LED), and pink is metal halide (MH). The distribution of lighting technologies along road transects represent the situation post 2014. Pre-2014 the lights shown as LED were HPS. Minor and major roads are shown as grey and black lines, respectively. The only major road in Jersey runs along the central-southern coast (long row of MH lights). Coloured dots in (C) represent bat passes for different species/species groups from all iBats car-driven surveys from 2011-2015 (n=110 recording events).

**Figure 2.**
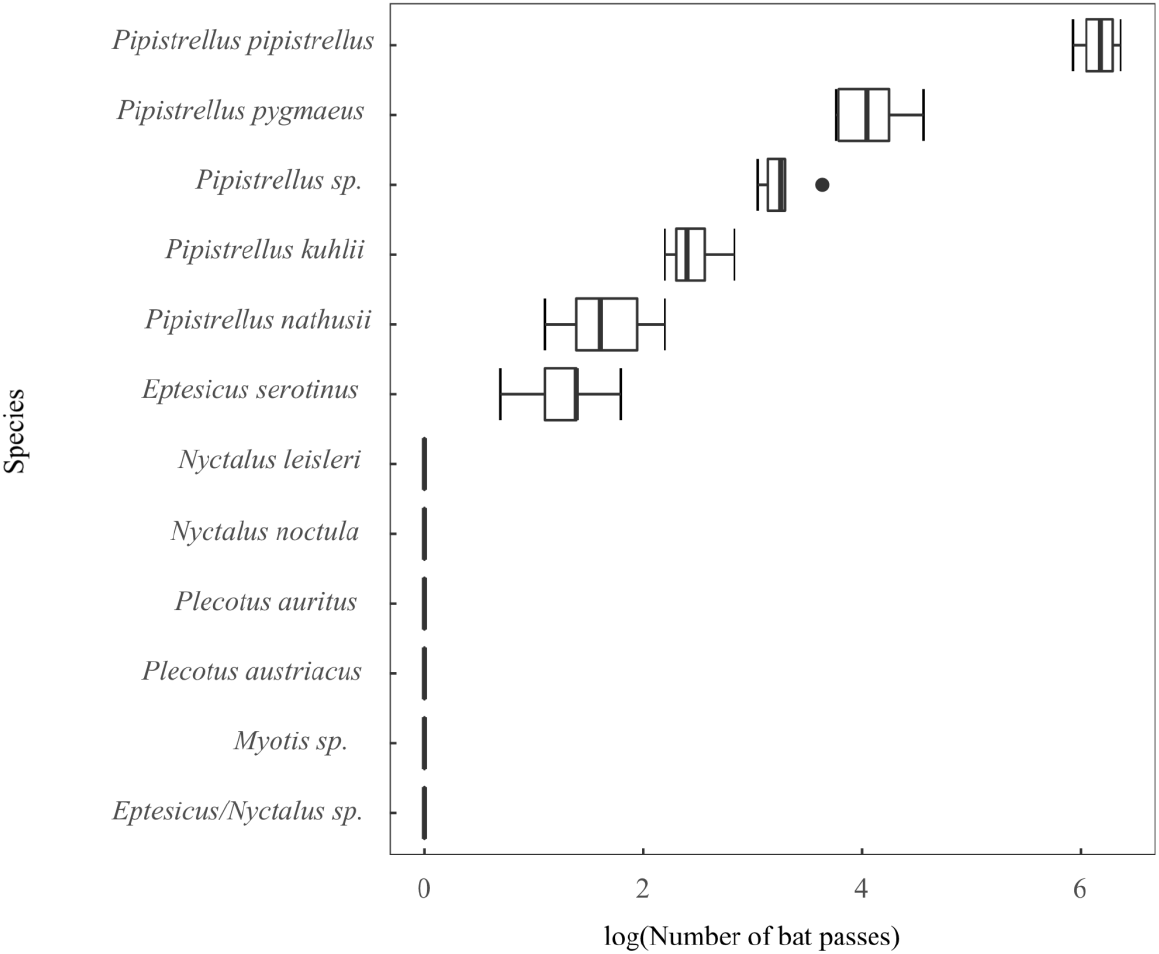
Number of bat passes recorded per iBats car-driven survey in Jersey from 2011-2015 for each species or species group, shown on natural log scale. Boxes show median and interquartile range and whiskers represent variability outside the upper and lower quartiles, with outliers plotted as individual points. See Table S2 for yearly totals.

**Figure 3.**
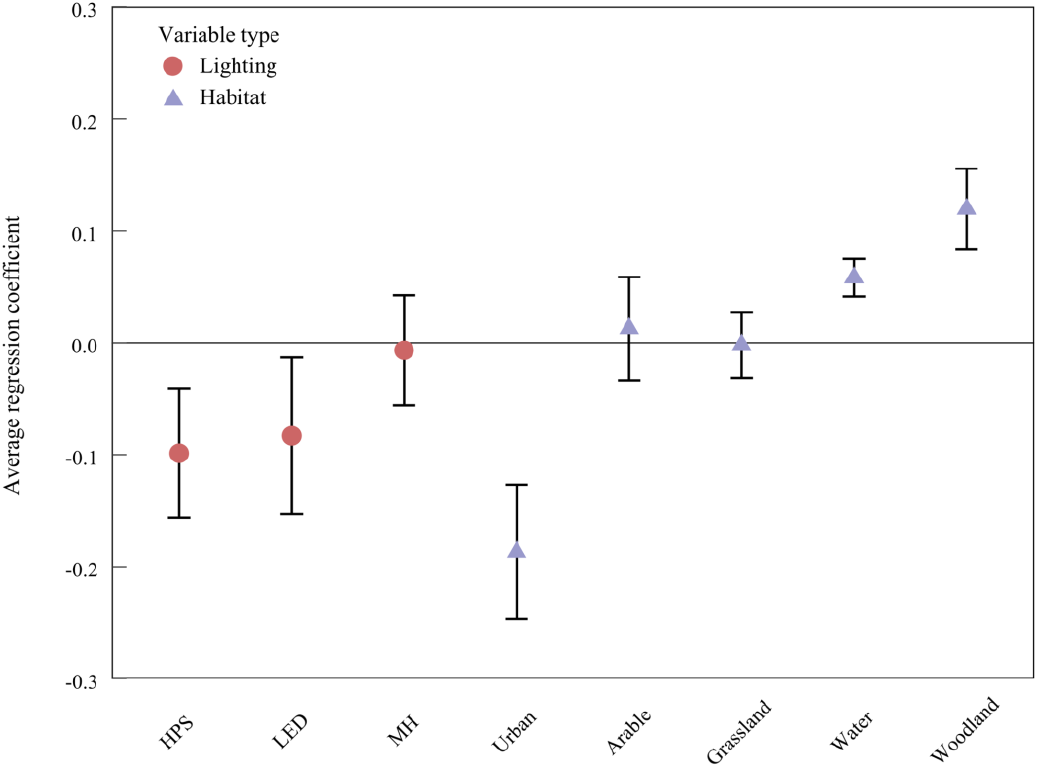
Average regression coefficients of lighting and habitat predictors from GLMM of *Pipstrelluspipistrellus.* Significant predictors are those whose 95% confidence interval (shown as error bars) does not include zero. The predictors (all lighting and habitat variables) shown have been centred and scaled (subtracting mean and dividing by standard deviation), meaning that the magnitude of the effects of each predictor variable on bat activity (the average regression coefficients) are directly comparable. Climatic and road-type variables are not shown, as they represented factors which could not be scaled, so coefficients could not be compared.

**Table 2:**
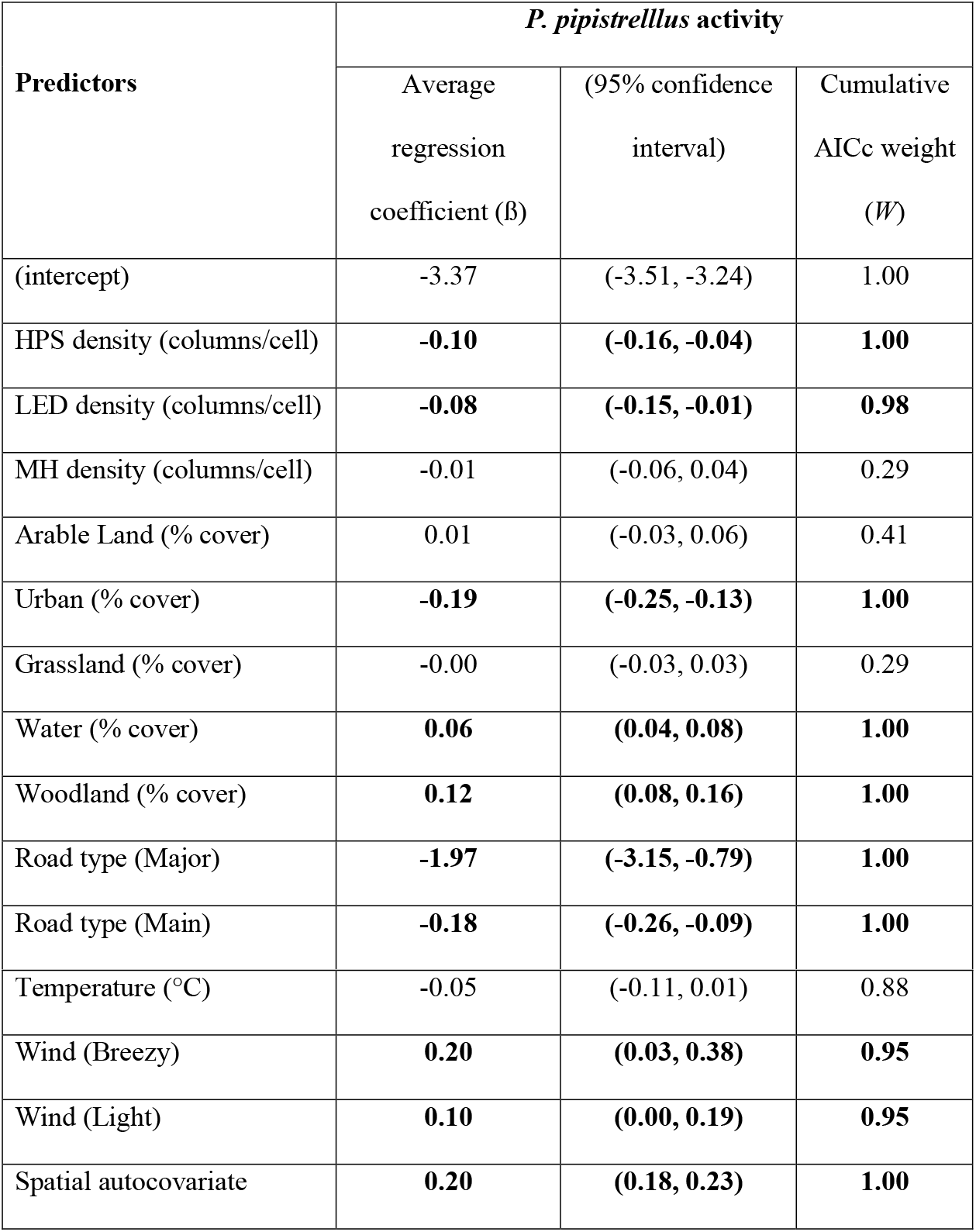
Averaged GLMM examining landscape effects of broad-spectrum lighting technologies and other spatial environmental variables on *Pipistrellus pipistrellus* activity. Regression coefficients (β) and 95% confidence intervals (,) are derived from each set of informative models (ΔAIC<7) (see Supplementary Information Table S3). *W* = cumulative AICc weights of all informative models that contain a given predictor variable. Significant predictors are shown in bold. All the predictor variables except road type and wind were centred and scaled (subtracting mean and dividing by standard deviation), meaning that the magnitude of each predictor variables effect on bat activity (the average regression coefficients) are directly comparable.

| Predictors | *P. pipistrellus* activity |  |  |
| --- | --- | --- | --- |
| | Average regression coefficient ($\beta$) | (95% confidence interval) | Cumulative AICc weight ($W$) |
| (intercept) | -3.37 | (-3.51, -3.24) | 1.00 |
| HPS density (columns/cell) | **-0.10** | **(-0.16, -0.04)** | **1.00** |
| LED density (columns/cell) | **-0.08** | **(-0.15, -0.01)** | **0.98** |
| MH density (columns/cell) | -0.01 | (-0.06, 0.04) | 0.29 |
| Arable Land (% cover) | 0.01 | (-0.03, 0.06) | 0.41 |
| Urban (% cover) | **-0.19** | **(-0.25, -0.13)** | **1.00** |
| Grassland (% cover) | -0.00 | (-0.03, 0.03) | 0.29 |
| Water (% cover) | **0.06** | **(0.04, 0.08)** | **1.00** |
| Woodland (% cover) | **0.12** | **(0.08, 0.16)** | **1.00** |
| Road type (Major) | **-1.97** | **(-3.15, -0.79)** | **1.00** |
| Road type (Main) | **-0.18** | **(-0.26, -0.09)** | **1.00** |
| Temperature (°C) | -0.05 | (-0.11, 0.01) | 0.88 |
| Wind (Breezy) | **0.20** | **(0.03, 0.38)** | **0.95** |
| Wind (Light) | **0.10** | **(0.00, 0.19)** | **0.95** |
| Spatial autocovariate | **0.20** | **(0.18, 0.23)** | **1.00** |

Our results were also robust to conducting the same analyses at a different spatial scale (100×100m grid cells; Table S4).

## Discussion

### i) Landscape effects of different broad spectrum lighting technologies

Our results reveal differences in the relative impacts of three different broad-spectrum streetlight technologies on the activity of a bat species, *P.pipistrellus*, that is often found in urban habitats and generally considered to be resilient to human disturbance (8). Whilst MH density had little impact on this species’ activity, the two non-UV emitting technologies LED and HPS had a significant and negative impact on activity, across multiple years of monitoring at a large (island-wide) spatial scale.

Previous studies found that experimental illumination of hedgerows with HPS and LED streetlights had no significant effect on *P. pipistrellus* activity (8,22). However, these were short and small in scale, and our findings for HPS density are supported by recent evidence from Ireland of lower *Pipistrellus* activity in road sections lit using HPS lights (23). Our study is the first to reveal a similarly negative interaction between LED lighting and landscape-scale *P. pipistrellus* activity, again despite previous experimental studies having suggested otherwise (and despite this species typically being characterised as light-tolerant). These apparent contradictions between experimental results and our larger-scale findings suggest that the spatiotemporal scale of analyses may substantially influence our ability to reliably detect the effects of artificial lighting on populations. This highlights the importance of conducting future studies at spatial scales that are most relevant to detecting population-level effects. We also advise caution around use of the term ‘light-tolerant’ in bat-lighting research, as our results alongside others (18,23) suggest that artificial lighting has the potential to detrimentally affect the ecology of many apparently light-tolerant species.

The differences in the average regression coefficients observed between the three light types may be related to UV content. Relative to MH lights, fewer insects should be attracted to HPS and LED lights, since these do not emit UV (19). A recent experimental study showed that significantly higher numbers of insects were attracted to MH light than to HPS and LED, which both attracted similar numbers of insects (28). There is also experimental evidence linking bat activity to streetlight UV content, for example significant decreases in *Pipistrellus* and *Nyctalus/Eptesicus sp.* activity after replacing high-UV with low-UV light (18,21). As such, the negative relationship we observe between HPS/LED density and *P.pipistrellus* activity may result from a combination of limited feeding opportunities and local increases in light-dependent predation risk around artificial lights (23,29). Conversely, increased predation risk around MH lights could be offset by local increases in foraging success, which may explain the relatively neutral relationship between MH density and *P.pipistrellus* activity.

Unlike previous experimental analyses, our study examined the effects of lights that bats will have become habituated to. We also accounted for temporal transitions from yellow HPS to white LED lights that occurred in certain parts of the study area from 2014 onward. However, the exact dates when each HPS light was replaced with an LED were unavailable, and it is possible that LEDs were not installed in certain areas until 2015 or later, which could confound the parameter estimates for these lighting types. Thus, care in interpreting our results concerning the effect of LED lights should be taken, but these estimates seem intuitive considering the well-established link between UV content and bat behavior.

By modelling key environmental and road-type variables alongside lighting densities, we were also able to control for habitat heterogeneity, degree of urbanisation, road-type and weather, all of which had noticeable effects on patterns of bat activity. In particular, intact woodland and aquatic habitats were consistently the strongest positive predictors of *P. pipistrellus* activity around roads, which are in accordance with other studies of bat space use, and likely relate to higher insect abundances in these habitat types (30,31). Roads act as barriers to bat movement (4), so efforts to improve permeability by improving roadside habitat are a priority in bat conservation; our results suggest that, as well as preserving woodland and wetland habitat, it is crucially important that artificial lighting regimes are, wherever possible, excluded from areas of critical habitat (Figure 4).

**Figure 4.**
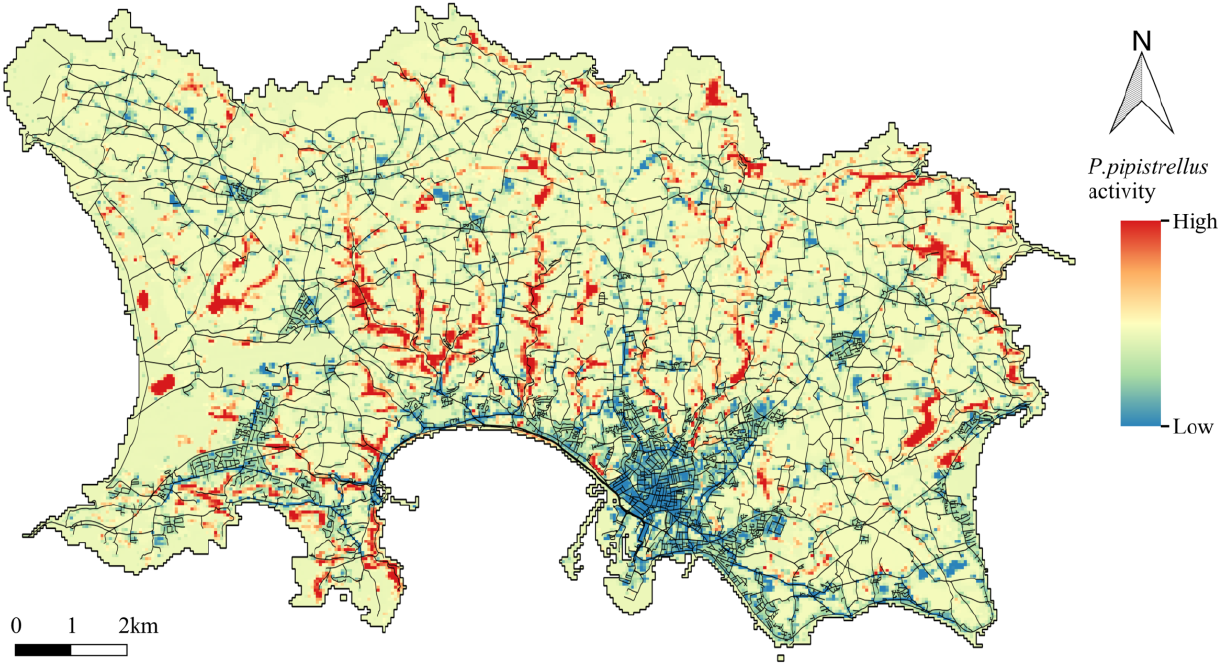
Spatial predictions of *P.pipistrellus* activity in the absence of artificial lighting from the averaged GLMM. Note that grid cell values are the result of prediction from an averaged model based in part on scaled variables so have no direct interpretation. Red corresponds to habitat where activity is predicted to be highest and blue, the lowest. Roads (black lines) are also shown to enable identification of the areas that may be particularly sensitive to placement of streetlights.

### ii) Future recommendations

Considering our findings that the two emerging technologies LED and MH have different effects on *P. pipistrellus* activity at an island-wide scale, we suggest that which of these light-types is chosen to replace end of life HPS lights could differentially impact the diversity and species composition of bat communities. Conversion of existing HPS lights to MH lights may provide generalist species such as *P.pipistrellus* with increased foraging success in lit environments. However, the high UV content of MH lights could lead to detrimental community level effects by increasing the ‘vacuum effect’ (28) and hence reducing food availability for genera such as *Plecotus* and *Rhinolophus* which tend to be excluded from illuminated areas (8,16,22). There is some evidence for lighting regimes causing such ecological displacements. For example, declines in *Rhinolophus hipposideros* populations in Switzerland may have been a result of competitive exclusion by *P. pipistrellus* (32).

Increased attraction of bats to MH lights could also exacerbate the effect of roads on bats by increasing the chance of vehicular collision. We therefore advise against the use of high UV content streetlights such as MH where possible, due to their potential to cause far-reaching ecological disturbance.

In the coming years, the States of Jersey plan to replace all streetlights with LED bulbs (Jersey Electricity Company pers. comm.). Considering the similarity in model estimates for HPS and LED lighting and their abilities to attract insects (28), conversion of existing HPS lights to LEDs may not noticeably affect bat communities. However, such a transition would involve a subtle change in light colour from yellow (HPS) to white (LED). Experimental research using lamps which emitted negligible amounts of UV showed that *P. pipstrellus* activity was significantly higher around white light than red light and no light (33). However, further research is needed into how changing light colour from yellow to white whilst holding UV content constant may impact upon bat behaviour. Recent experimental evidence showed that activity of *Myotis spp.* (a light-averse genus) increased significantly when high UV mercury vapour lights were converted to LED lights, whilst activity of *P. pipistrellus* was significantly reduced. Thus, it was suggested that LED lights could potentially benefit light-averse species by reducing the barrier effects of high UV lighting and by reducing any potential competitive advantage of light-tolerant species (18). Despite the negative effect of LED observed here on *P. pipistrellus*, proposed replacement of old lamps with LEDs could potentially reduce long-term anthropogenic disturbance to bat communities by minimizing the disparity between the effects of artificial lighting on generalist species (such as *P. pipistrellus)* and on those that are more obviously light-averse (such as *Myotis* spp.).

Installation of lighting regimes along roads has the potential to exacerbate their barrier effects and create vacuum effects for insect biomass. Thus, it is important that lights are only installed in key areas, away from areas of critical habitat (Figure 4). Whilst such an approach should be implemented where possible, we also recognize the role that artificial lighting regimes may play in increasing the safety of roads. Advances in streetlight technologies mean that there are now numerous possible solutions to minimize ecological impact. For example, directed lighting and motion sensors could be used to restrict unnecessary light usage. The UV content of streetlight technologies is also non-functional for humans, so removal of UV wavelengths could have far-reaching ecological benefits without compromising performance (23). Due to the inherent taxonomic biases in road-based acoustic surveys, we were unable to examine the effect of each streetlight technology on species with a greater light intolerance. Given that road artificial lighting regimes are increasing and energy efficient technologies such as LED are increasing in popularity, this should also become a research priority. Nonetheless, our results stress that artificial lighting regimes have complex landscape-scale effects on bat ecology and distributions, even for species broadly considered to be human-tolerant. Our study also adds to a growing body of evidence showing the impressive ability of citizen science data to reliably detect global change effects on species ecology, such as the negative impacts of artificial light. Innovative citizen science programmes will likely play an increasingly important role in long-term monitoring of biodiversity.

## Material and Methods

### (i) Bat acoustic activity data

Bat echolocation calls were collected as part of the Indicator Bats (iBats) Programme conducted on the island of Jersey (49.2144° N, 2.1312° W) (Figure 1). Ultrasonic recordings were made along 11 transect routes of approximately 25 km, repeated twice (and occasionally three times) in July and August of each year from 2011-2015, using full-spectrum time-expansion (TE) acoustic devices (Tranquility Transect, Courtplan Design Ltd, UK), following the methods in (26) (see Supplementary Information Methods for detailed data collection protocols). Each single survey event is henceforth referred to as a *‘recording event’.* Transects were driven at 25 km/hr starting 30-45 minutes after sunset, and a GPS track was simultaneously recorded along the transect route to enable geo-referencing of recorded calls. At the beginning of each recording event, data on temperature (° C) and wind speed (calm, light or breezy) were recorded (surveys were not conducted in high winds). Acoustic devices were set to record using a TE factor of 10, a sampling time of 320ms, and sensitivity set on maximum, giving a continuous sequence of ‘snapshots’, consisting of 320ms of silence (sensor listening) and 3.2 seconds of audio (sensor playback at x 10 speed). Sound was recorded to an SD card as a WAV file using either an Edirol R-09HR or Roland R-05 recording device. Due to occasional equipment failure or unforeseen circumstances, eight recording events were slightly modified before analysis or deleted, resulting in 110 recording events (total of 132.1 hours) from 11 transects (Figure 1a) (see Supplementary Information Table S1 for details).

Audio from each recording event was split into one minute segments and the presence of bat echolocation calls was manually verified visually using Batsound software (Pettersson Elektronik). Segments containing echolocation calls were individually detected, and high-quality calls were located and call parameters were extracted using Sonobat v.3.1.7p (34). Parameterisation in Sonobat involves the use of amplitude threshold filters and recognition of smooth frequency changes over time to find calls and to fit a frequency-time trend line to the shape of the call, from which measurements are extracted. An artificial neural network (ANN) for European species, iBatsID (25) was used to classify calls to species level, however calls with a classification probability of under 60% were only classified to species-group level, as either; *‘Pipistrellus\* ‘Serotine/Leisler’s/Noctule’, or ‘Long-eared’. Calls identified to any *Myotis* species were reclassified as ‘*Myotis*’ due to the difficulty of distinguishing these species. For *P.pipistrellus*, the focal species of this study, iBatsID has a correct call classification rate of 97.6% and a false positive rate of 1.4% (25). Individual calls were grouped into sequences *(‘batpasses’)* for analysis, where calls were assumed to be part of the same call sequence if they occurred within the same snapshot and if the sequence continued into subsequent snapshots. Each sequence was classified to a species or species-group using either the species or species-group class (if species classifications were equivocal) of the majority of the echolocation calls (Figure 1b). If there is no majority species ID to which the sequence can be assigned, the sequence is assigned to the group stage (e.g. ‘Unknown *Pipistrellus*’). We divided the island of Jersey into 50 x 50m grid cells and calculated the total number of bat passes from each species in each 50 x 50m grid cell that intersected a transect road per recording event. This spatial resolution was used as it was assumed to be the maximum size at which only one sampled road was included in each grid square.

### (ii) Street lighting technology and environmental spatial data

Streetlights were located along bat transect roads using data compiled from the States of Jersey lighting surveys conducted in 2011 and updated in 2017. As only approximate locations were provided, we manually ground-truthed the exact position of all streetlights along the transect roads. Data on model type and wattage of each streetlight from the Jersey Electricity Company (JEC) was used to determine the locations of lights belonging to the three different technologies: high pressure sodium (HPS), metal halide (MH), and light emitting diode (LED) (Figure 1c). In 2014, the States of Jersey began to replace end of life HPS streetlights with LEDs. We created two datasets, pre and post-2014, to account for these changes. The pre-2014 dataset includes only HPS (n=935) and MH streetlights (n=138). The post-2014 dataset includes HPS (n=599), MH (n=138) and LED streetlights (n=336)) accounting for any changes in HPS streetlights to LEDs that occurred from 2014 until the latest lighting survey in 2017. Data concerning the exact date from 2014-2017 when each end of life HPS light was replaced with LEDs was unavailable, so we assume an immediate transition (i.e. all LED lights recorded in the lighting survey of 2017 were present from 2014 onwards). We then created 50 x 50m resolution lighting layers for our analysis, displaying the number of columns of each streetlight type in each cell pre and post-2014. As some roads in Jersey are not lit by streetlights, our analysis inherently compares bat activity in unlit areas versus lit areas.

We also collected information on habitat and road-type variables found to be important correlates of bat activity (4,35,36). Data on six key habitat variables (arable, grassland, water, woodland, urban and unclassified) were extracted from a Phase 1 Habitat Survey of Jersey in 2011, provided by the States of Jersey (37) (Supplementary Information Figure S1). The original shapefile was converted into a 2m^2^ resolution gridded layer for each habitat type using the *raster* package (38) in R (39). We chose this resolution to ensure maximum preservation of spatial information whilst maintaining computational efficiency. Percentage cover of each habitat type was calculated for each 50 x 50m grid cell which intersected a bat transect road by aggregating the 2m2 grid cells in the high-resolution raster. Bat transect roads were categorised into three types using the definitions in the Phase 1 Habitat Survey; ‘Minor’, ‘Main’ and ‘Major’ (Figure 1C), and we created a 50 x 50m resolution grid of all cells that intersected transect roads, whose values corresponded to the type of road in each cell. In the few cases that multiple road types occurred per cell, the road type that covered the greatest area was assigned. We also created separate 50 x 50m resolution grids displaying the temperature and wind speed values recorded at the start of each recording event.

### (iii) Statistical analyses

We used a generalised linear mixed model (GLMM) with a Poisson error distribution using the lme4 package in R version 3.4.2 (39) to examine the relationship between *P.pipistrellus* activity and densities of different streetlight technologies, habitat and road-type, and climatic variables in 50 x 50m grids on and around bat transect roads. We also included a temporal random effect in the model, where month was nested within year, as we found a temporal trend of increasing bat activity over the study period (2011-2015) (Supplementary Information Figure S2, Table S2) and to account for the repeated sampling of each grid cell. Transect ID was not included as a random effect because transects exhibited minimal spatial overlap and were each surveyed only once per month, and therefore a month nested within year random effect is sufficient to account for repeated sampling of grid cells. The *P.pipistrellus* activity data were slightly zero-inflated which could impact parameter estimates, however we found little difference in results when fitting a zero-inflated Poisson model in glmmTMB package in R version 3.4.2 (39), so we do not report the results here. Bat activity data recorded from 2011-13 and from 2014-2015 were paired with the pre-and post-2014 lighting technology datasets, respectively. Continuous predictors (all lighting density and habitat-type variables plus temperature) were centred and scaled (subtracting mean and dividing by standard deviation) to facilitate model convergence and to make their effect sizes biologically interpretable (40). To account for spatial autocorrelation, an autocovariate was included as an additional predictor variable (41). This is calculated for each 50 x 50m grid cell as a distance-weighted average of neighbouring response values (number of detections per cell) (42). Neighbourhood distance was set to the minimum distance at which no cells had zero neighbours (spdep package in R version 3.4.2) (43,44). All variables were checked for multicollinearity by calculating correlation coefficients between each variable. In all cases, r < 0.7 so no variables were excluded (45).

We adopted an information-theoretic approach (IT) to construct averaged models using the*MuMIn* package (46). Firstly, all the variables were used in the initial model. Next, a comprehensive set of models was derived that represented every permutation of all the variables specified in the initial model. The corrected small-sample Akaike Information Criterion weight (AICc) and the difference between a given model and the model with the lowest AIC (ΔAIC) was then calculated for each model. Any model with a ΔAIC >7 was deemed uninformative and discarded (47). The average parameter estimate of each variable was then calculated across all informative models, and its relative importance was estimated by summing the AICc weights of all informative models that included that particular variable (27) (see Supplementary Information Table S3 for all informative models). Since the results of our spatial analysis could be sensitive to spatial scale, we also conducted these analyses at a grid cell size of 100 x 100m. The results for both scales were qualitatively similar (Supplementary Information Table S4), so we present our models here using 50 x 50m grid cells.

The averaged model of *P.pipistrellus* activity constructed at 50 x 50m was then projected across the entire island of Jersey using the percentage cover data from each habitat variable in all cells (including those in which no sampled road was found). This provided a visual representation of the spatial variation in bat activity and enabled the identification of critical habitat areas that may be particularly sensitive to placement of lighting. As data on lighting variables were only available for the sampled cells, they were removed from the predictive process by setting the island wide-density per cell to zero (i.e. predicting habitat suitability in the absence of lighting). Island-wide data on variables that represented factors (wind and road-type) were also unavailable for non-sampled cells. To enable prediction, they were set to their base condition (‘Calm’ for wind, ‘Minor’ for road-type in all cells (including sampled cells).

## Data Accessibility

All code and data is available on figshare:

1. Code: https://figshare.com/s/340bfb9e7530698d9818
2. Raw data: https://figshare.com/s/c29d41f47f0a10299721
3. Processed data: https://figshare.com/s/a1b5745966559428596d

## Acknowledgements

We thank H Ferguson-Gow for technical assistance, and Department for Infrastructure, States of Jersey and the Jersey Electricity Company (JEC) for providing lighting data. This work was funded with support from States of Jersey (KEJ and CW), a Graduate Research Scholarship from University College London (RG), and NERC PhD studentship (NE/L002485/1) (EB).

## Supplementary Information Methods

### Indicator Bats Programme Protocols

Citizen scientists within the iBats programme (Jones et al. 2013) collected full-spectrum acoustic recordings of approximately 70 minutes, along ~25km road transects routes, driven at 25 km/h and starting 30-45 minutes after sunset. Each transect was surveyed twice yearly during July to August (the period of peak seasonal activity for most northern European species) spanning a period from 2011-2015 across the island of Jersey. Acoustic data was recorded with a time-expansion (TE) device equipped with a wideband capacitance microphone with the frequency range 10-160 kHz and a sample rate of 409.6 kHz (Tranquility Transect, Courtplan Design Ltd, UK). The recording devices were set to record using a TE factor of 10, sampling time of 320 ms and sensitivity set on maximum, giving a continuous sequence of ‘snapshots’, consisting of 320 ms of silence (sensor listening) and 3.2s of TE audio played backed 10 times slower than real time. The recording device was attached to the front or back passenger window of the vehicle with the microphone pointing up and back at a 45° angle. Edirol R-09HR or Roland R-05 recording devices were used to record audio without compression or using a lossless compression, and native files were transformed into wav format. A GPS track of the transect route was recorded using a Samsung GT S77710 phone with a built in Global Positioning System (GPS) taking readings every 5 seconds via OruxMaps or other GPS options, and the transect tracks were recorded as GPX files. The sound recorder and GPS track were set to start recording at the same time so that the position of the car when each bat is recorded could be determined. Each recorded bat call was subsequently georeferenced using this GPS track. Surveys were only carried out during ‘fine’ weather; i.e. when the air temperature was greater than 7°C, and no more than very light rain or wind. The following metadata were also recorded at the start and end of each transect: temperature (°C); cloud cover (%); rain (dry, drizzle, light); and wind speed (calm, light, or breezy). Humidity data for the start and end of each transect was obtained from the States of Jersey Department for Environment - Meteorological Section.

**Table S1.**
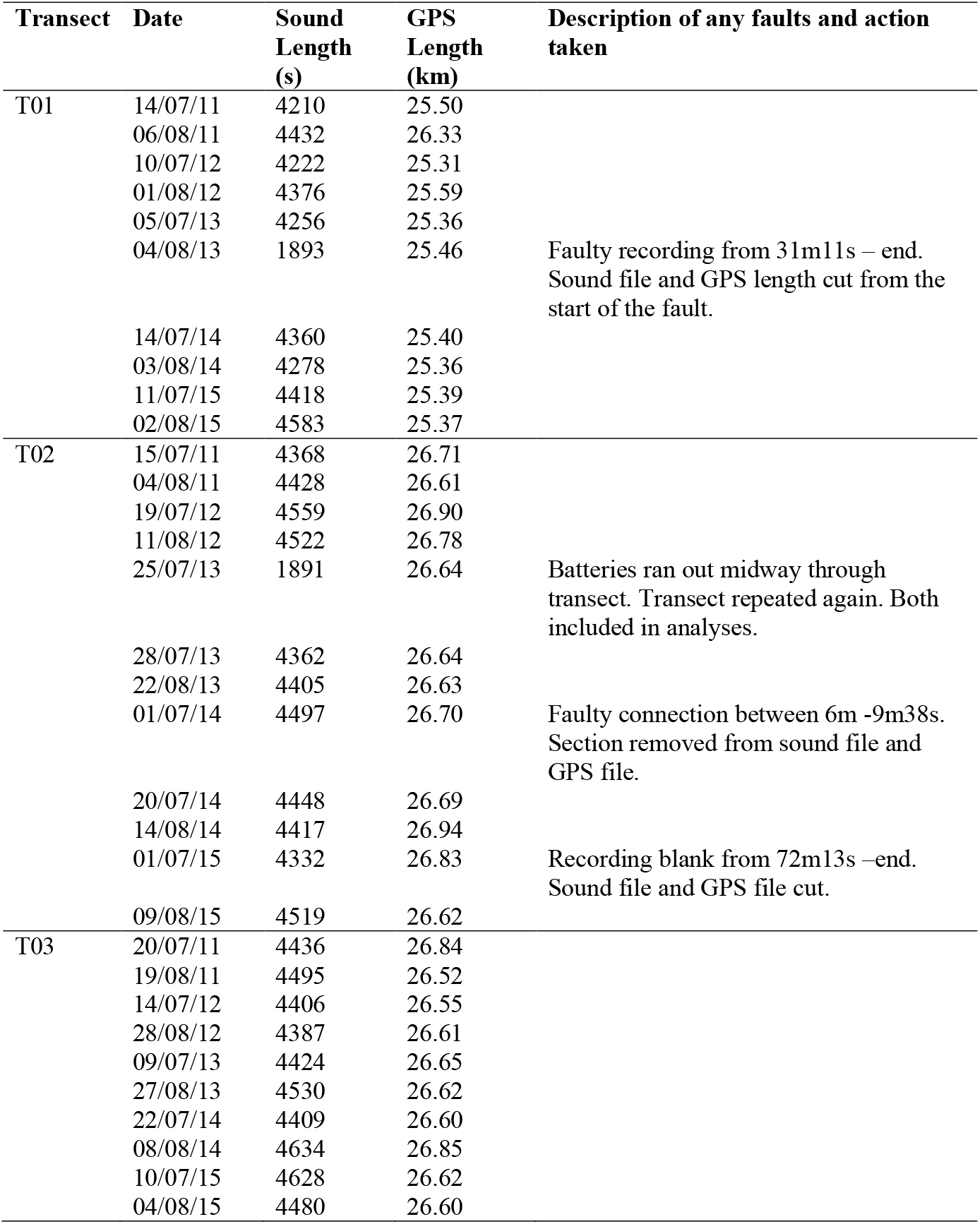
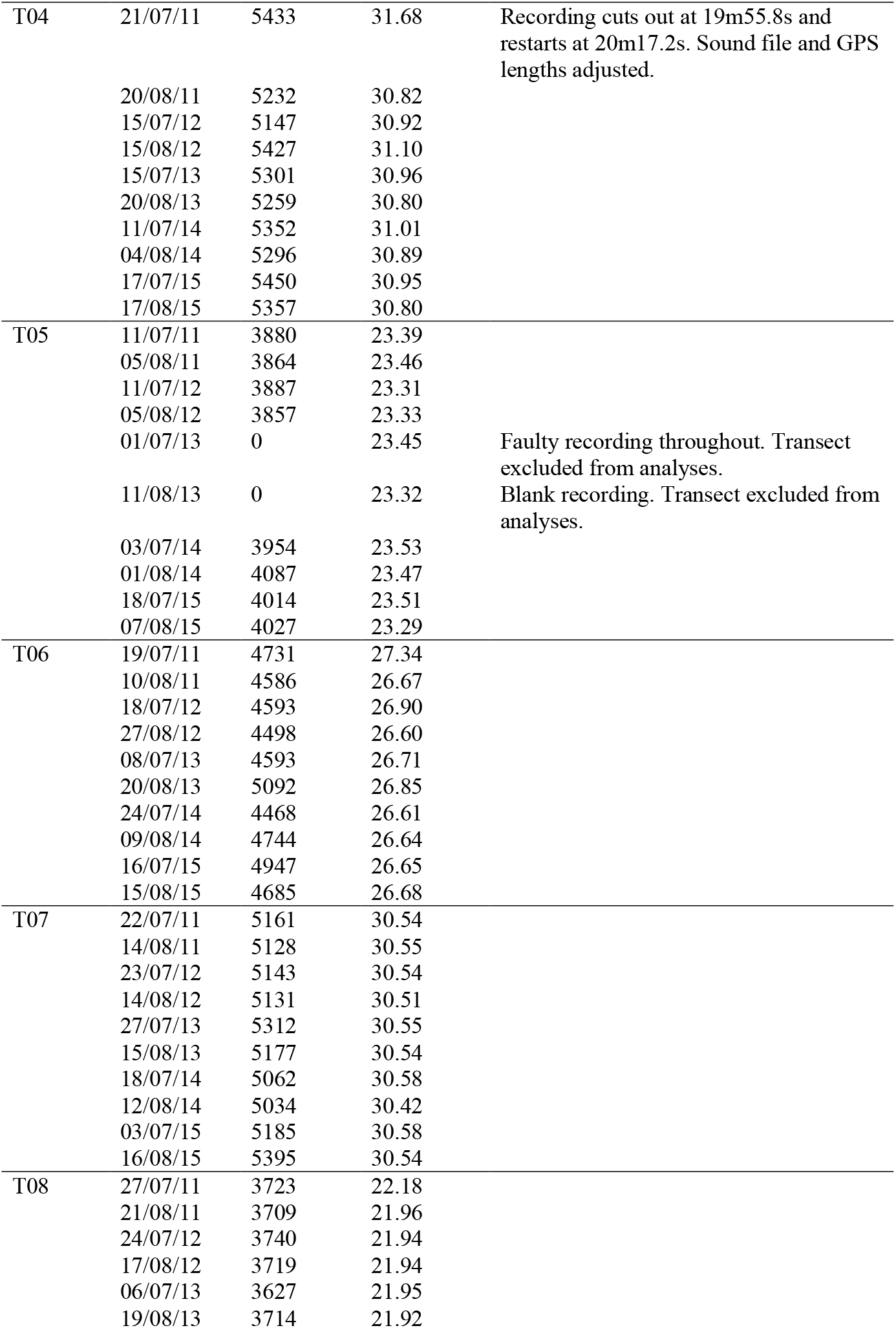
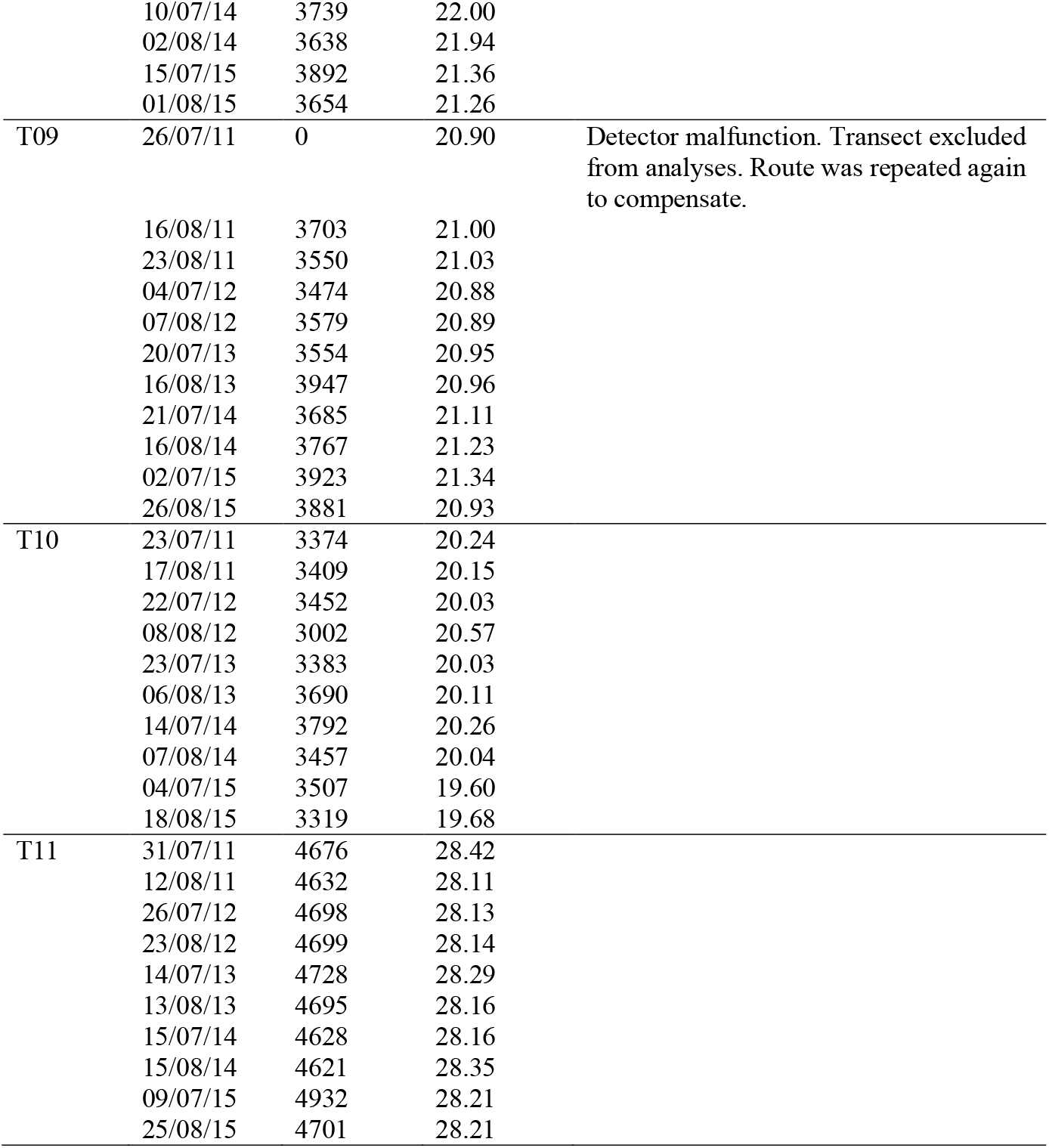
Details of the iBats car driven acoustic transects (n=11) carried out between 2011-2015 on the island of Jersey, including the faulty recordings and how these were used in the analysis.

**Figure S1.**
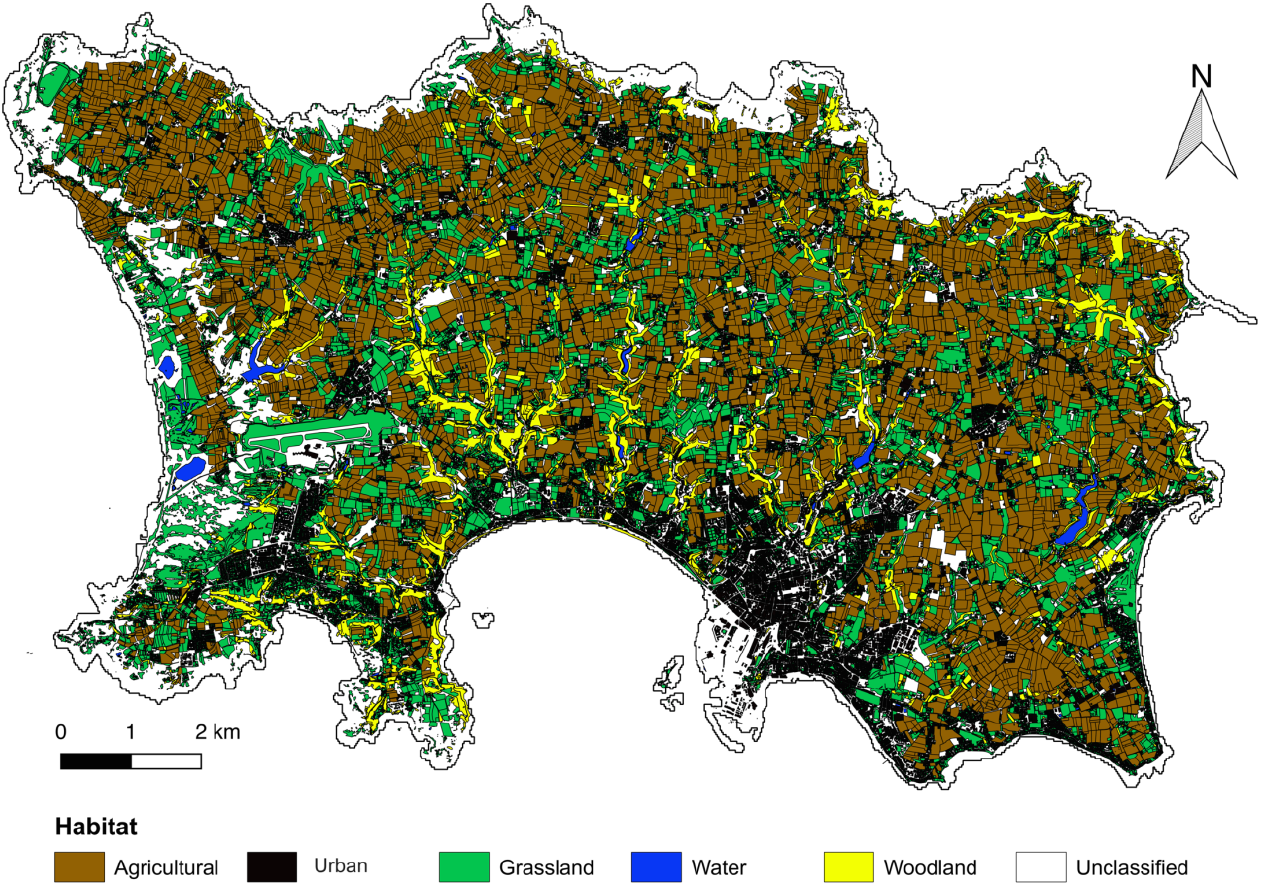
Distribution of six key habitat types across the island of Jersey. Habitats were grouped using the Phase 1 Habitat Survey classes of Jersey are follows: (1) Arable land, including habitats defined as arable land and arable land short term ley; (2) Urban, using habitats defined as buildings; (3) Grassland including areas defined as improved, semi-improved and unimproved grassland, marshy (improved, semi-improved, unimproved and Oenanthe dominated) grassland, coastal (Molinia, dune and species rich short turf) grassland, amenity/parkland, gardened, and saltmarsh; (4) Water, including areas defined as standing and running water, brackish pool, swamp; (5) Woodland, including areas defined as plantation, planted broadleaved, semi-natural broadleaved, coniferous, deciduous and mixed woodland; and (6) Unclassified.

**Figure S2.**
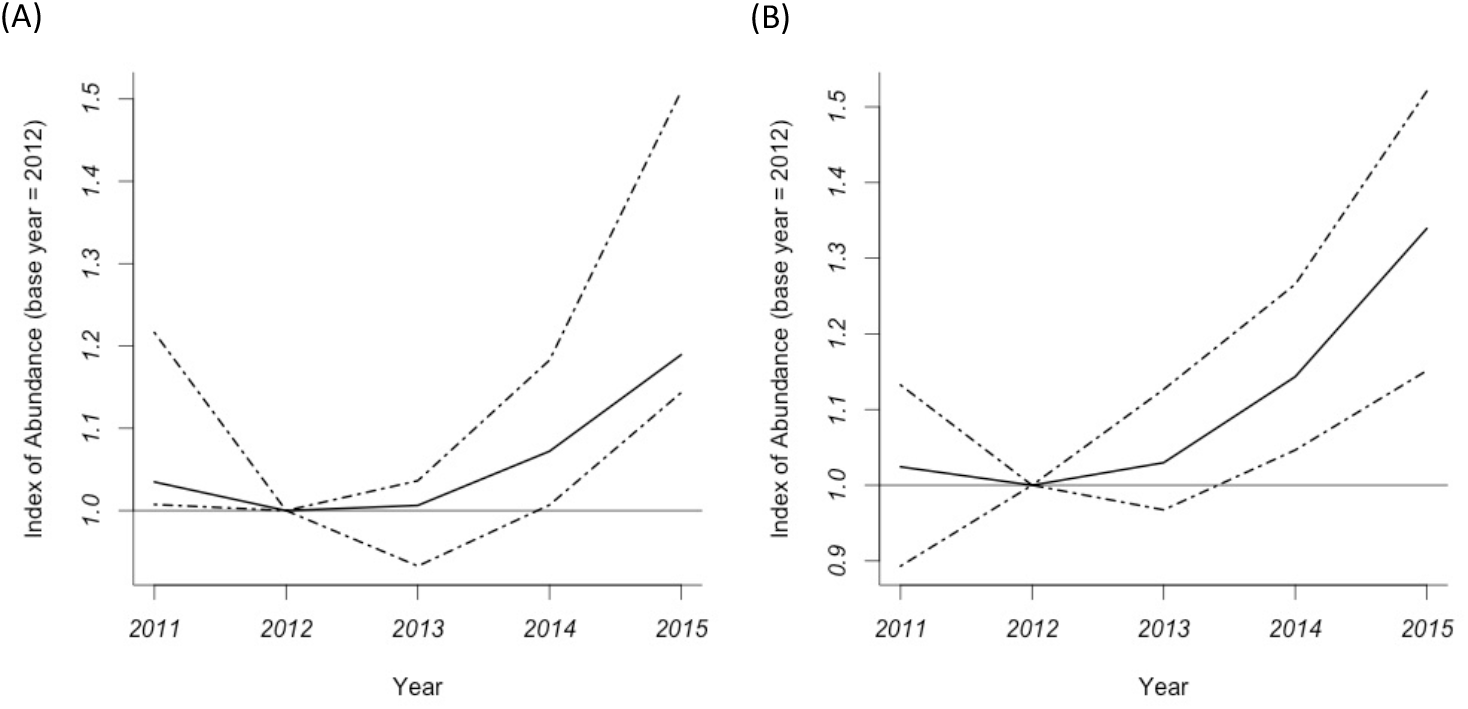
Population trends of (A) total bat passes and for (B) the common pipistrelle *(Pipistrellus pipistrellus)* only across Jersey from 2011-2015. Population trends were estimated using a Generalized Additive Model (GAM) with a Poisson error distribution following (Barlow et al., 2015; Fewster et al., 2000). The baseline year (index = 1.0) was set as the second year in the time series (2012). Degrees of smoothing were set to 0.4 times the number of years of survey data. Confidence intervals were obtained by bootstrapping directly from the index curve. 399 bootstrap replicates were obtained and the confidence intervals were set to 95%. The significance of the trend (at the 5% significance level) was determined by whether the confidence intervals in the final year of surveys (2015) overlapped with the population abundance index of the baseline year (2012). Passes for the measure of total activity were identified using a species-level classification probability threshold of zero, and percentage cloud cover was included as a covariate in the trend analyses. Start temperature and start cloud cover were included as covariates in trend analysis for *P. pipistrellus*. GAMs were constructed using mgcv 1.8-4 in R (Wood 2011). Total bat activity and *P. pipistrellus* activity significantly increased by approximately 19% and 34% between 2012-2015, respectively.

**Table S2.** Number of passes of each bat species or species group recorded per year in Jersey with the percentage of total annual bat passes in parentheses.

| Species | Number of passes (% of total annual bat passes) |  |  |  |  |
| --- | --- | --- | --- | --- | --- |
|  | 2011 | 2012 | 2013 | 2014 | 2015 |
| *Pipistrellus pipistrellus* | 484<br>(83.45) | 426<br>(80.38) | 376<br>(81.39) | 540<br>(77.47) | 582<br>(82.55) |
| *Pipistrellus pygmaeus* | 57<br>(9.83) | 44<br>(8.30) | 43<br>(9.31) | 96<br>(13.77) | 70<br>(9.93) |
| *Pipistrellus kuhlii* | 9<br>(1.55) | 17<br>(3.21) | 10<br>(2.16) | 13<br>(1.87) | 11<br>(1.56) |
| *Pipistrellus nathusii* | 3<br>(0.52) | 9<br>(1.70) | 5<br>(1.08) | 4<br>(0.57) | 7<br>(0.99) |
| *Nyctalus leisleri* | 0<br>(0) | 1<br>(0.19) | 1<br>(0.22) | 1<br>(0.14) | 0<br>(0) |
| *Nyctalus noctula* | 1<br>(0.17) | 1<br>(0.19) | 0<br>(0) | 0<br>(0) | 1<br>(0.14) |
| *Eptesicus serotinus* | 4<br>(0.69) | 4<br>(0.75) | 2<br>(0.43) | 3<br>(0.43) | 6<br>(0.85) |
| *Plecotus auritus* | 0<br>(0) | 0<br>(0) | 1<br>(0.22) | 0<br>(0) | 0<br>(0) |
| *Plecotus austriacus* | 0<br>(0) | 0<br>(0) | 0<br>(0) | 1<br>(0.14) | 1<br>(0.14) |
| *Pipistrellus sp.* | 21<br>(3.62) | 27<br>(5.09) | 23(4.98) | 38<br>(5.45) | 26<br>(3.69) |
| *Eptesicus/Nyctalus sp.* | 1<br>(0.17) | 1<br>(0.19) | 0<br>(0) | 1<br>(0.14) | 1<br>(0.14) |
| *Myotis sp.* | 0<br>(0) | 0<br>(0) | 1<br>(0.22) | 0<br>(0) | 0<br>(0) |
| **TOTAL Species** | **6** | **7** | **8** | **7** | **7** |
| **TOTAL Passes** | **580** | **530** | **462** | **697** | **705** |

**Table S3.**
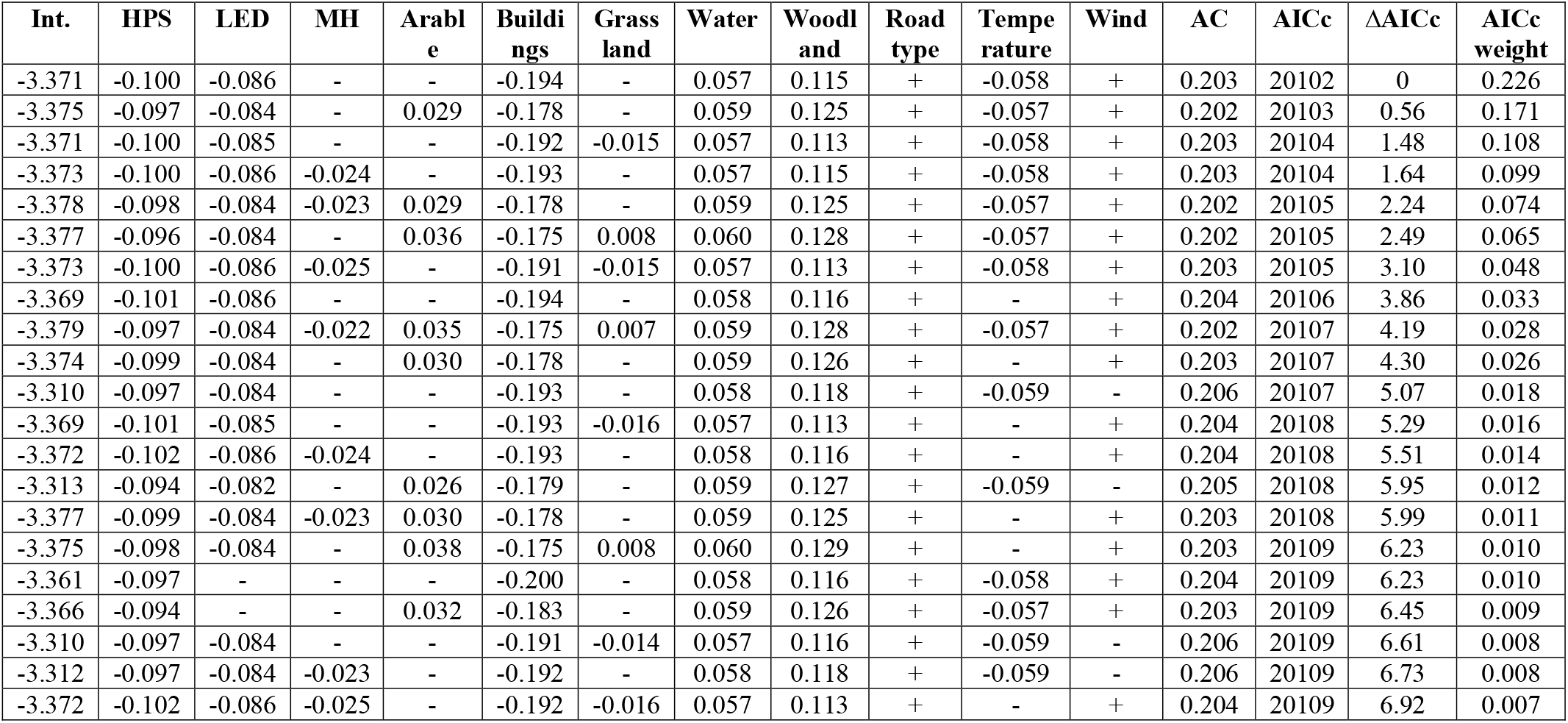
All informative models (ΔAICc < 7) used to construct average GLMM of *P.pipistrellus* activity (50 x 50m). AC = autocovariate, △AIC = the difference between a given model and the model with the lowest AICc, AIC weight = cumulative AICc weights of all informative models that contain a given predictor variable.

**Table S4.** All informative models (ΔAICc < 7) used to construct average GLMM of *P.pipistrellus* activity (100 x 100m). AC = autocovariate, △AIC = the difference between a given model and the model with the lowest AICc, AIC weight = cumulative AICc weights of all informative models that contain a given predictor variable.

| Int. | AC | Arable | Buildings | Grassland | HPS | LED | MH | Road type | Temperature | Water | Wind | Woodland | AICc | $\Delta\text{AICc}$ | AICc weight |
| --- | --- | --- | --- | --- | --- | --- | --- | --- | --- | --- | --- | --- | --- | --- | --- |
| -2.679 | 0.188 | 0.092 | -0.128 | 0.046 | -0.098 | -0.069 | - | + | -0.045 | 0.060 | + | 0.192 | 16933 | 0 | 0.169 |
| -2.672 | 0.189 | 0.049 | -0.144 | - | -0.102 | -0.071 | - | + | -0.045 | 0.055 | + | 0.172 | 16933 | 0.06 | 0.165 |
| -2.665 | 0.190 | - | -0.171 | - | -0.105 | -0.073 | - | + | -0.046 | 0.052 | + | 0.154 | 16934 | 1.53 | 0.079 |
| -2.678 | 0.188 | 0.093 | -0.128 | 0.046 | -0.099 | -0.069 | - | + | - | 0.060 | + | 0.193 | 16934 | 1.62 | 0.075 |
| -2.671 | 0.190 | 0.050 | -0.144 | - | -0.103 | -0.071 | - | + | - | 0.055 | + | 0.173 | 16934 | 1.66 | 0.074 |
| -2.678 | 0.188 | 0.092 | -0.128 | 0.046 | -0.098 | -0.069 | 0.005 | + | -0.045 | 0.060 | + | 0.192 | 16935 | 1.99 | 0.063 |
| -2.672 | 0.189 | 0.049 | -0.144 | - | -0.102 | -0.071 | 0.001 | + | -0.045 | 0.055 | + | 0.172 | 16935 | 2.06 | 0.061 |
| -2.664 | 0.191 | - | -0.172 | - | -0.106 | -0.073 | - | + | - | 0.052 | + | 0.155 | 16936 | 3.29 | 0.033 |
| -2.665 | 0.190 | - | -0.169 | -0.010 | -0.105 | -0.073 | - | + | -0.046 | 0.051 | + | 0.153 | 16936 | 3.32 | 0.032 |
| -2.665 | 0.190 | - | -0.171 | - | -0.105 | -0.073 | 0.000 | + | -0.046 | 0.052 | + | 0.154 | 16936 | 3.53 | 0.029 |
| -2.677 | 0.188 | 0.093 | -0.128 | 0.046 | -0.099 | -0.069 | 0.005 | + | - | 0.060 | + | 0.193 | 16936 | 3.61 | 0.028 |
| -2.671 | 0.190 | 0.050 | -0.144 | - | -0.103 | -0.071 | 0.002 | + | - | 0.055 | + | 0.173 | 16936 | 3.66 | 0.027 |
| -2.671 | 0.189 | 0.098 | -0.133 | 0.049 | -0.094 | - | - | + | -0.045 | 0.060 | + | 0.195 | 16936 | 3.68 | 0.027 |
| -2.664 | 0.190 | 0.051 | -0.151 | - | -0.098 | - | - | + | -0.045 | 0.055 | + | 0.173 | 16937 | 4.00 | 0.023 |
| -2.664 | 0.191 | - | -0.170 | -0.010 | -0.106 | -0.073 | - | + | - | 0.051 | + | 0.154 | 16938 | 5.05 | 0.014 |
| -2.664 | 0.191 | - | -0.172 | - | -0.106 | -0.073 | 0.001 | + | - | 0.052 | + | 0.155 | 16938 | 5.29 | 0.012 |
| -2.670 | 0.189 | 0.099 | -0.133 | 0.049 | -0.095 | - | - | + | - | 0.060 | + | 0.196 | 16938 | 5.30 | 0.012 |
| -2.665 | 0.190 | - | -0.169 | -0.010 | -0.105 | -0.073 | 0.000 | + | -0.046 | 0.051 | + | 0.153 | 16938 | 5.32 | 0.012 |
| -2.608 | 0.192 | 0.045 | -0.145 | - | -0.099 | -0.069 | - | + | -0.046 | 0.055 | - | 0.173 | 16938 | 5.40 | 0.011 |
| -2.663 | 0.191 | 0.053 | -0.151 | - | -0.099 | - | - | + | - | 0.055 | + | 0.174 | 16938 | 5.59 | 0.010 |
| -2.613 | 0.191 | 0.086 | -0.130 | 0.043 | -0.095 | -0.067 | - | + | -0.047 | 0.060 | - | 0.192 | 16938 | 5.60 | 0.010 |
| -2.670 | 0.189 | 0.098 | -0.134 | 0.049 | -0.094 | - | 0.007 | + | -0.045 | 0.060 | + | 0.195 | 16938 | 5.65 | 0.010 |
| -2.656 | 0.192 | - | -0.180 | - | -0.101 | - | - | + | -0.046 | 0.052 | + | 0.155 | 16939 | 5.89 | 0.009 |
| -2.664 | 0.190 | 0.052 | -0.151 | - | -0.098 | - | 0.004 | + | -0.045 | 0.055 | + | 0.173 | 16939 | 5.99 | 0.008 |
| -2.602 | 0.194 | - | -0.170 | - | -0.102 | -0.071 | - | + | -0.047 | 0.052 | - | 0.157 | 16939 | 6.36 | 0.007 |

